# KaryoScan: abnormal karyotype detection from whole-exome sequence

**DOI:** 10.1101/204719

**Authors:** Evan K. Maxwell, Claudia Gonzaga-Jauregui, Shane E. McCarthy, Colm O’Dushlaine, Jeffrey Staples, Alexander E. Lopez, Xiaodong Bai, John Penn, Rick Ulloa, Omri Gottesman, Aris Baras, Frederick E. Dewey, Jeffrey Staples, John D. Overton, Erik Puffenberger, Lukas Habegger, Jeffrey G. Reid

**Affiliations:** Regeneron Genetics Center, Regeneron Pharmaceuticals, Tarrytown NY.; The Clinic for Special Children, Strasburg, PA.

## Abstract

**Motivation:** Detection of abnormal karyotypes from whole-exome sequencing has significant clinical potential, enabling a primary screen for chromosomal anomalies among samples undergoing short-read sequencing for nucleotide resolution genomic characterization.

**Results:** We present KaryoScan, a high-throughput method for detecting chromosomal anomalies within large cohort exome sequencing studies. We detect and validate autosomal and sex chromosomal aneuploidies in a large exome sequencing cohort, and demonstrate detection of smaller and complex events (partial chromosome, mosaic, copy neutral, and complex rearrangements), representing the range of anomalies that can be uncovered from the exome.

**Availability:** https://github.com/rgcgithub/karyoscan

## 1 Introduction

Karyotype assessment has traditionally involved chromosomal staining techniques and, more recently, array-based technologies, but the growing use of next-generation sequencing (NGS) as a primary diagnostic technology in both clinical and research settings has motivated the need for methods to detect karyotypes from NGS data. Particularly in large-scale, precision medicine initiatives aimed at screening patient DNA for pathogenic mutations, the ability to detect chromosomal anomalies without a secondary screening platform has significant advantages. Large array-based studies have shown that chromosomal anomalies are more frequent than might be expected from neonatal-based rates among adult clinical populations, particularly due to increased rates of mosaicism (both due to increased rates of hematological cancers as well as age-related clonal mosaicism) (Laurie *et al.*, 2012; Jacobs *et al.*, 2012). While methods to detect targeted classes of chromosomal anomalies (e.g trisomy 21, 13, and 18) from whole-genome sequencing (WGS) have been developed within the contexts of non-invasive prenatal testing (Ehrich *et al.*, 2011; Fan *et al.*, 2008; Chu *et al.*, 2009; Sehnert *et al.*, 2011), high-throughput methods for analyzing whole-exome sequence (WES) data for chromosomal anomalies in samples ascertained through standard clinical and research settings, where the primary goal is to assess nucleotide resolution variation, have not been developed.

Some complex karyotypes cannot be easily dissected with NGS approaches (e.g. ring chromosomes, isochromosomes, and very low-frequency mosaic anomalies); WES and targeted sequening panels are further limited by the presumed absence of breakpoint-spanning sequence necessary to detect genomic rearrangement events like deletions, duplications, inversions, and translocations. Thus WES-based karyotyping is focused on the detection of anomalies affecting chromosomal dosage (copy number) and allelic imbalance (e.g. uniparental disomy), with levels of detectable mosaicism being a function of the sequencing depth. In this manuscript, we present KaryoScan; a method for high-throughput detection of chromosomal anomalies from whole-exome sequencing.

## 2 Methods

**Fig. 1.**
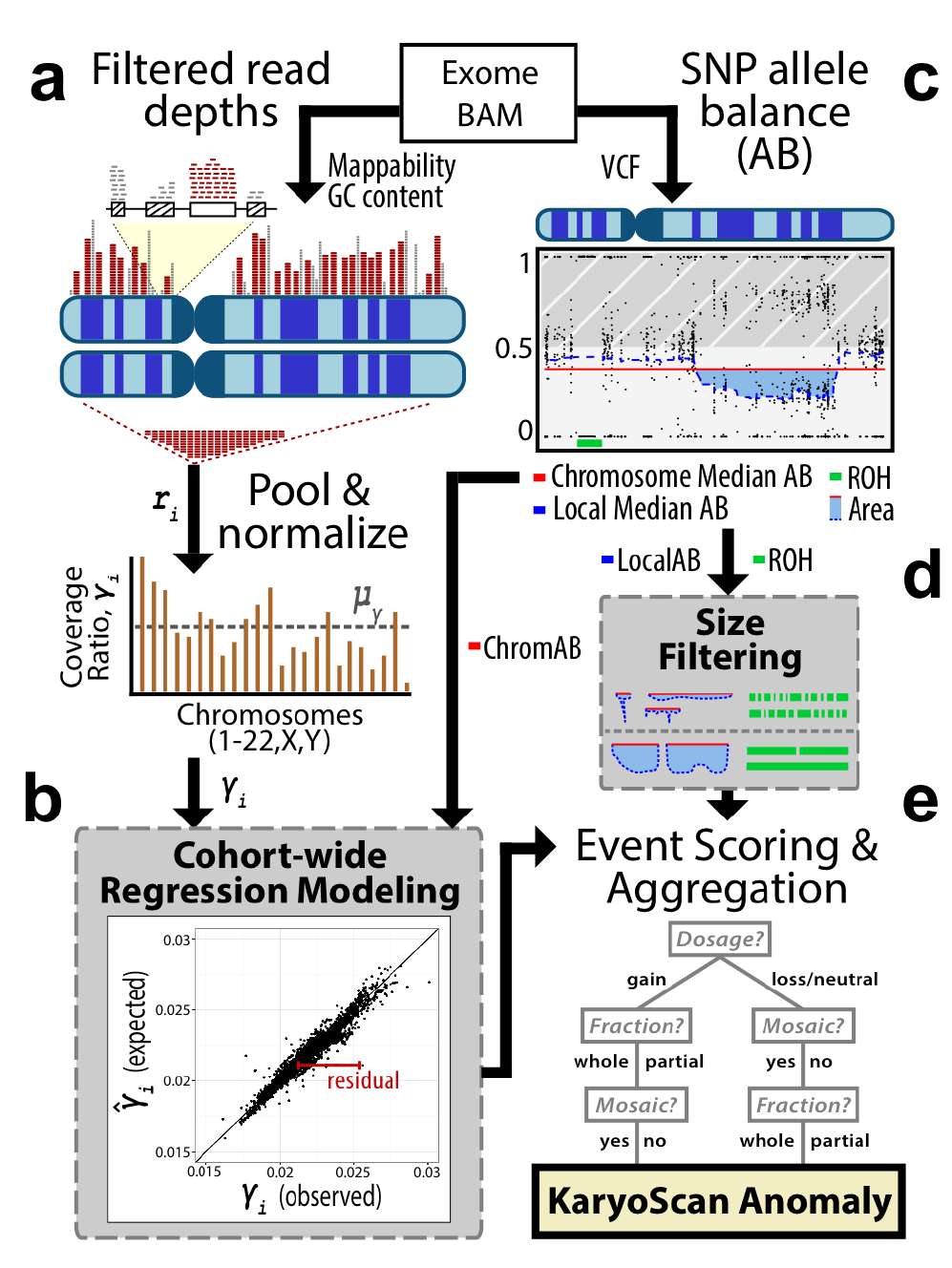
Overview of the KaryoScan algorithm. Depth-of-coverage and VCF files are computed from BAM files. **(A)** Read depths from high-confidence exonic regions are pooled across chromosomes, normalized, and **(B)** fit to a regression model to identify anomalous signals. **(C)** Biallelic SNPs from VCF files are analyzed based on allele balance (AB) to identify large runs-of-homozygosity (ROH) and chromosome-wide or partial chromosome (“local”) regions of anomalous heterozygous AB (differing from ~50%). Determination of anomalous allele balance signals are either **(B)** modelled cohort-wide or **(D)** identified by size filtering. **(E)** Signals from read depths and allele balance are scored and aggregated to produce a final karyotype anomaly call from KaryoScan.

### Chromosomal dosage assessment via read depths (RD)

Several technical artifacts obfuscate the assessment of sequencing read depths as an estimate of underlying genomic DNA dosage, including PCR amplification bias relative to GC content. We previously developed scalable normalization strategies for high-resolution detection of exonic copy-number variants (CNVs) with CLAMMS (Packer *et al.*, 2016); we apply and expand upon those principles within the KaryoScan framework. Briefly, CLAMMS partitions the exome into exonic windows, where a window represents either an entire exon or a contiguous 500–1000bp sub-segment for exons longer than 1kbp. Windows are then annotated for GC content, mappability, and known problem regions (e.g. pseudogenes, multicopy duplications, high mutational rates).

For detecting chromosomal anomalies within KaryoScan, we first compute a robust read depth profile r_i_ for each chromosome *i* as the sum of read depths over exon windows that are highly mappable (>95%), outside of problem regions, and have 45–55% GC content (Figure 1A). This strategy is better suited for detecting partial-chromosome gains and losses than the median chromosomal tag density metric used by some existing whole-chromosome aneuploidy detection methods (Chu *et al.*, 2009; Fan *et al.*, 2008) because the median may be unaffected or have a non-linear relationship with a partial-chromosome event. We then define the normalized coverage ratio γ_i_ of chromosome *i* as γ_i_=*r_i_*/Σ_*j*_r_*j*_, for all autosomes *j* s.t. *i ≠ j.* Notably, we compute *γ*_*i*_ for all chromosomes (autosomes and sex chromosomes), but normalize only to total autosomal read depth (removing the effect of sex) with the tested autosome excluded (to avoid dampening the signal of a trisomy, for example). We compare the ratios between chrX and chrY to define genetic sex and assess duplications of chrY (**Figure S1**).

Dosage altering chromosomal anomalies manifest in deviations of γ_i_ from expectation. However, the expected value of γ*_i_*(*E*[γ_i_])of is not constant even between karyotypically normal (diploid) samples. For every autosome and chrX, we fit a linear regression to predict *E*[γ_i_] for every individual (Figure 1B), with covariates from seven sequencing QC metrics (**Supplmental Methods**), the ratios of the two most similar chromosomes in terms of GC content, and, for chrX, genetically assigned sex (**Figures S1–2**). Using the fit model, we assess the difference between the observed and expected ratio (residual) for each individual on each chromosome to estimate the chromosomal gain or loss, as well as the significance of the deviation (Z-score transformation) to distinguish true chromosomal anomalies from noise (**Supplemental Methods**).

### Allelic imbalance assessment via heterozygous allele balance (AB)

Chromosomal anomalies can also be identified through analysis of allele balance (the fraction of reads supporting each allele) across a contiguous chromosomal range of single-nucleotide polymorphisms (SNPs). For example, a trisomy generally occurs through a duplication of one chromosomal copy, yielding heterozygous SNPs with allele balance fractions converging on 1/3 and 2/3. Similarly, a nonmosaic monosomy will be absent of heterozygosity. In the “normal” diploid state, heterozygosity is expected to occur at relatively consistent rates over the genome with allele balance modeled as a binomial distribution with mean=50% and variance a function of sequencing depth; allele balance should converge on 50% when summarized across all heterozygous positions on a chromosome (**Figure S3**). However, to detect allele balance anomalies (i.e. regions of allelic imbalance where the heterozygous allele balance ≠ 50%), we focus only on the “lesser allele balance” (henceforth denoted “AB”): the fraction of reads supporting the less frequent allele, defined in the range of [0–0.5] (Figure 1C). Due to this selection criteria, AB approaches 0.5 as sequencing depth increases, but will converge on a value <0.5 (**Figure S3**, **Supplemental Methods**).

To detect abnormal karyotypes from allele balance, KaryoScan parses single-sample VCF files, computing (**Supplemental Methods**):

1) Chromosome-wide median AB value for all high-quality, putative heterozygous SNPs (“ChromHetAB”)
2) Local median AB value for all high-quality, putative heterozygous SNPs, using a running median over a 20 SNP window (“Lo-calHetAB”)
3) Runs-of-homozygosity (“ROH”): contiguous regions absent of hets, computed with BCFtools/RoH (Narasimhan *et al.*, 2016)

Anomalous values are detected independently for each metric (Figure 1B & D). For ChromHetAB, we model the expected value for each chromosome relative to a representative exome-wide coverage metric (we use percent target bases covered at 50х or greater, **Figure S4**), trained across the entire sample cohort (Figure 1B). Anomalous values are based on thresholds for Z-score transformed residuals using a one-tailed test. For LocalHetAB, we identify chromosomal segments where the LocalHetAB value is lower than the ChromHetAB value over a series of heterozygous SNPs, and compute a normalized “area under the curve,” for each event, filtering events with area < 0.02 (**Figure 1D & S6**). Similarly for ROH segments, we filter events having fewer than 40 SNPs and spanning less than 5 Mbp.

### Aggregation of signals

Following acquisition of a set of putative chromosomal anomalies stemming from independent read depth and allele balance metrics (Figure 1B & D), we normalize scores from different metrics to a common “tier” scale. We define tiers 1–3 for each metric, where tier 1 events are the most significant, and tier 3 are the least (**Supplemental Methods**). All putative anomalies are then aggregated within a particular chromosome for a particular individual (Figure 1E) to determine a final, chromosome specific call containing an overall tier rating as well as predictions of dosage change (or copy neutral), whole or partial chromosome, mosaic or not, and estimates of the chromosomal and/or mosaic fraction detected. Samples having at least one chromosomal anomaly flagged are assigned to tiers equal to their most significant event’s chromosomal tier, and diagnostic plots showing read depth distributions and allele balance across the exome are generated for every flagged sample, enabling manual inspection of karyotype calls.

## 3 Results

We trained KaryoScan on the Regeneron Genetics Center’s human exome variant database samples, and tested our method’s sensitivity and specificity on a set of 2,084 samples from the Clinic for Special Children (CSC). The core set of 1,698 samples consist of pedigrees associated with probands referred for genetic testing and their family members. The majority of the probands (N=116) in this core set have undergone clinical grade CytoScan testing for diagnosis, and DNA from family members for which a KaryoScan anomaly is detected are available for CytoScan testing. This core set provides a readily available data set for orthogonal validation of KaryoScan. An additional 386 samples are derived from older cell lines where immortalizaion in culture may have enabled expansion of anomalous clones; we only attempted validation of these KaryoScan anomalies if the event was assumed to be germline.

Of note, autozygosity rates are known to be high among Amish and Mennonite populations ranging from 2.5% to 4.1% (Strauss and Puffenberger, 2009). We detected 561 samples having ROH segments encompassing at least 25% of a chromosome with KaryoScan, however no samples had whole chromosome UPD events detected. Of the 2,084 CSC samples, KaryoScan identified two trisomy 21 cases and four (47,XXY) male cases in the core sample set, as well as one (45,XO) female among the non-core sample set, all scored as tier 1 dosage-altering anomalies (**Figures S7A-13A**). Both trisomy 21 cases have a clinical diagnosis of Down Syndrome and were previously confirmed via microsatellite genotyping. None of the sex chromosome anomalies were known *a priori,* and all five were subsequently confirmed with CytoScan (**Figures S7B-13B**). KaryoScan also identified two core samples having mosaic copy neutral events of subchromosome scale on chromosomes 8 and 11, neither of which were known clinically and both validated with CytoScan (**Figures S27–28**). The mosaic nature of these events suggest they were acquired (i.e. somatic), differentiating them from the regions of autozygosity frequently identified within the population. No false negative chromosomal anomalies were identified among probands with existing CytoScan data (N=98). In a separate study, we identified large CNVs (>50 exons; e.g. 16p13.11, 22q11.2, 1q21. 1, 15q11.2) in 27 core samples with CLAMMS (Packer *et al.*, 2016); these carriers were correctly deemed karyotypically normal with KaryoScan.

Among the 386 non-core samples, several tier 1 complex karyotypes were identified by KaryoScan including three mosaic monosomy X cases (46,XX/45,XO), one mosaic trisomy X (46,XX/47,XXX), a putative non-mosaic unbalanced translocation t(+1q,–16q), and eight additional mosaic duplication and deletion events detected on chromosomes 9, 12, 5q, 6p, 9q, 11q, 14q, 22q (**Supplemental Methods**). Review of the KaryoScan diagnostic plots (**Figures S14–26**) and support for the smaller events from existing CLAMMS CNV data suggest they are all true anomalies likely stemming from cell line artifacts. Together, these events demonstrate the range of detectable chromosomal anomalies from KaryoScan.

## Acknowledgements

This work was funded by the Regeneron Genetics Center.

